# Pen-strep influence macrophage mechanical property and mechano-response to microenvironment

**DOI:** 10.1101/2020.04.09.034884

**Authors:** Weikang Zhao, Buwei Hu, Xuexiang Zhang, Pingping Wang

## Abstract

Penicillin-streptomycin (Pen-strep) is a common antibiotic used to prevent bacterial infection in cell culture and clinical treatment. Current research found pen-strep increased macrophage modulus but limited influence on cell adhesion. Phalloidin statin image indicates pen-strep mediate cell morphology on different extracellular matrix coated surface. The roundness analyzes further illustrated pen-strep promote cell spread on PDMS rubber, type I collagen, laminin, poly amino acid, poly-RGD peptides. Finally, YAP-1 and TAZ upregulation and β1 integrin downregulation may be the causes of cell elasticity and mechano-response to extracellular matrix (ECM) change.

## Introduction

Macrophage derived from monocyte and participates in several immune responses including defense of bacterial infection, regulates inflammation and tissue repairment [1-3]. During immune response, macrophage will further convert into M1 and M2 subtypes. M1 is pro-inflammatory phase, which secretes TNF-α, IL-12, and other cytokines to initiate immune cascade. The M2 phenotype has anti-inflammatory properties, it can repress immune response and promote extracellular matrix (ECM) reorganization, tissue regeneration [4].

Cell elasticity is an important characteristic which reflects cell physiological condition. More specifically, macrophage cell modulus could indicate macrophage subtype and function [5, 6]. Through LPS stimulation, the elasticity of activated macrophage is three times higher than resting macrophage. This research also indicated that activated macrophage increases its adhesion strength [7]. Furthermore, compared with M2, M1 macrophage has higher cell plasticity [6]. Base on this discovery, macrophage modulus also found to influence cell phagocytosis, migration, and cytokines including reactive oxygen species (ROS) secretion [8]. Currently, study on chemical induced macrophage modulus change is mainly focused on LPS stimulation condition. The influence given by commonly used drug still needs further investigation.

Penicillin and streptomycin (pen-strep) are broad-spectrum antibiotics widely used in bacterial infection treatment and cell culture study [9, 10]. Our research found that pen-strep treatment influences macrophage plasticity and its sensation to microenvironment. In general, Pen-step increases macrophage modulus. It also further mediates cell sensation to extracellular matrix (ECM) and cell morphology. The analyze of protein expression indicate that YAP-1 and TAZ mediate cell modulus, Furthermore, the downregulation of β1 integrin may influence cell attach to different ECM.

## Method

### 1. Reagents

The reagents listed are required: AlexaFluor 488 Phalloidin (Thermo Fisher, A12379), Fluormount-G (SouthernBiotech, 0100-01), Triton X-100 (Sigma, T8787), 10X Phosphate Buffered Saline (PBS, Corning, 46-013-CM), 16% Paraformaldehyde (PFA, Electron Microscopy Science, 15710). Normal Serums and Gamma Globulins (NDS, Donkey, Jackson Immuno Research Labs, 017000121), 4’,6-Diamidino-2-Phenylindole (DAPI, Introgen™ Molecular Probes™, D3571)

### 2. Equipment and software

The equipment listed or similar alternatives are required: 25×75 mm Microscope slides (Fisher Scientific company, 12-550-15), 12 mm Microscope cover slip (Fisher Scientific company, 12CIR-1.5), Zeiss Fluorescence Microscope. ImageJ (https://imagej.nih.gov/ij/download.html) was used to measure cell roundness. AFM, Single cell

### 3. Reagent setup

10X PBS was stored at room temperature (20-25°C) and was diluted in distilled water to 1X working solution. 4% PFA was prepared from 16% PFA diluted in 10X PBS and distilled water. NDS stock solution was dissolved in 1X PBS and further diluted to 5% (w/v) NDS in 1X PBS. Triton X-100 was dissolved in 1X PBS to 0.5% (v/v) concentration. DAP I was diluted by PBS (1/1000) before staining and preserved in 4°C.

### 4. Extracellular matrix coating and PDMS prepare

We coated all the glass sheets with different materials. Each material(poly-RGD peptide, PDMS rubber, collagen 1, collagen 4, poly amino acid, laminin) is made into a 10mg /ml liquid under sterile conditions. The prepared coating materials will be put on the glass at 37C for half an hour.

### 5. Cell culture

RAW cells were obtained from the American Type Culture Collection (Manassas, VA) and cultivated in advanced Dulbecco’s modified Eagle’s medium (Invitrogen) with 10% FBS (Fisher Scientific, Houston, TX). And seeded into 24-well multiplates at 5,000 cells/well and used at 80% confluence. Two hundred microliters of culture media containing different concentrations (1–100 μg/mL) of PS was added directly to the cells, and the plates were incubated at 37°C for 24 hours in a humidified incubator with 95% air and 5% CO2. To determine the importance of physical contact between PS and cells for Cell mechanicals, the PS were suspended in the culture media (indirect exposure) in a separate experiment.

### 6. Polymerase chain reaction (PCR)

RAW cells were cocultured with small disks of different groups with diameters of 3 cm and thicknesses of 0.5 cm in a 6-well plate. DMEM/10% FBS was added to the cultured cells, and the cell-attachment-related and cell-modulus-related genes (TAZ, Egr-1, YAP-1, vinculin, paxillin and β1 integrin) were detected. The first established detection time point was 1 day after coculture with the samples.

RNA was extracted using a total RNA kit (Omega Bio-Tek, Norcross, GA, USA) at each experimental time point. Complementary DNA was synthesized (at 37 °C for 15 min and 85 °C for 5 s) using a PrimeScript RT reagent kit (Takara, Shiga, Japan). A quantitative reverse transcription-polymerase chain reaction (qRT-PCR) assay was performed using the SYBR premix Ex Taq reagent (Takara) with a CFX Connect Real-Time PCR Detection System (Bio-Rad, Hercules, CA, USA). Primers were designed and synthesized by Sangon Biotech Co. Ltd. (Shanghai, People’s Republic of China) using Primer Premier software (PREMIER Biosoft, Palo Alto, CA, USA). The primer sequences are presented in Table 1. Glyceraldehyde-3-phosphate dehydrogenase was used as an internal control. The relative expression of the target gene was calculated using the 2−ΔΔCt method according to the Ct values measured in previous reports.

### 7. Atomic force microscope and single cell force spectroscopy

A micromanipulator (Narishige International, Tokyo, Japan) was used to attach RAW to a “V” shaped tipless AFM cantilever (Bruker, NP-O10, Camarillo, CA, USA) under a microscope (Bruker, Camarillo, CA, USA) All AFM measurements were carried out at room temperature in sterile PBS (pH 6.0) using an optical lever microscope (Nanoscope V, Digital Instruments, Woodbury, NY, USA) with z-scan rates of 1.0 Hz under a maximal loading force of 5 nN. All cantilevers were calibrated using AFM Tune-it Version 2 software to get the spring constant of the cantilever. Force–Distance curves were measured at different contact time (0, 5 and 10 s). All force curves were analyzed using Nanoscope Analysis (Bruker, version 1.40) software. For every test a freshly prepared probe was used, and on each sample, force-distance curves were recorded thrice at three random spots and averaged.

### 8. Phalloidin cell skeleton fluorescence staining

Cell seed on glass slides or different ECM coating fixed by 4% PFA for 10 minutes. Then, washed by 1X PBS 3 times and penetrate by 0.5% triton X-100 in PBS for 20 min. Samples are incubate at room temperature for 2 h with 5% NDS for blocking. After that, samples are incubate with alexafluor 488 labelled phalloidin (1:400) for 1 hour in room temperature. Finally, samples were cultured with DAPI for 5 min. Between these steps, slides were washed with PBS three times for 5 min to remove unbonded molecules. Samples were mounted by fluormount media and sealed by microscope cover glass. Then slides will be put in 4°C refrigerator overnight to allow cover glass set before imaging. A zeiss microscope was used to image macrophage morphology. imageJ was used to measure the roundness of macrophage.

## Results

### Pen-strep increase macrophage modulus but no effect on adhesion

Macrophage elasticity indicates its function and influence its cytokine secretion [6]. Through atomic force microscope (AFM) measurement, cell stiffness can be detected base on the force applied on the indenter tip of cantilever (Figure. 1a). Cells were cultured on glass slides and treat with pen-strep for five days. Cellular stiffness was measured every day. Results showed that under pen-strep treatment, macrophage modulus increase after 24 hours and reach the highest modulus on day 5 at around 2.5 KPa (Figure. 1b). Furthermore, cell elasticity has a dramatic increase from day 2 to day 3, it raised from 1.5 to 2.3 KPa in 24 hours. In contrast to the influence on cellular modulus, cell focal adhesion does not have significant change under pen-strep treatment. Similar to AFM, the single-cell force spectroscopy measure cell adhesion force base on the force apply on cell to detach from substrate (Figure. 1c). After five days of culture and measurement, results indicate cell focal adhesion only be affected after two-day culture, reach the bottom point at around 5 nN. It back to normal, around 6 nN, and does not have significant change after day 3 (Figure. 1d).

**Figure 1:**
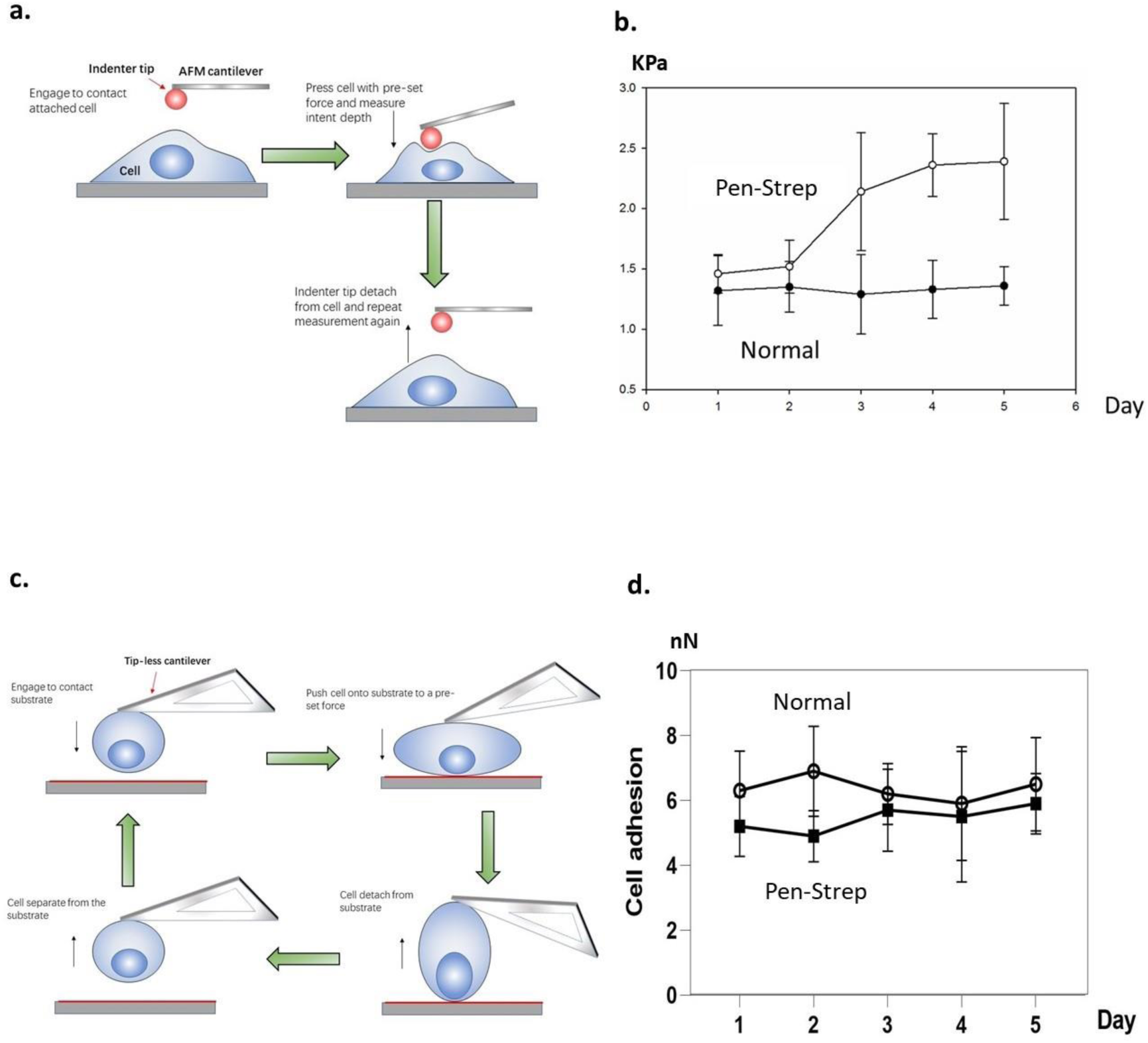
a) The illustration of atomic force microscopy mechanism; b) Change of cell modulus with or without Pen-Strep treatment for 5-day treatment, Unit: KPa; c) The illustration of single-cell force spectroscopy mechanism; d) Change of cell adhesion with or without Pen-Strep treatment for 5-day treatment, Unit: nN.

### Pen-strep influence macrophage microenvironment sensation

Having the observation on the change of cell elasticity after treatment, we hypothesis that pen-strep treatment influences cell morphology and even sensation to ECM. After seed cells on different materials or ECM treated substrate for 24 hours, phalloidin staining used to shown macrophage shape. Roundness used to indicate cell shape. After 24 hours treatment, the roundness of cell on glass slides does not have significant change compare with control group (Figure. 2a). Compare with glass, macrophage on PDMS rubber does not prefer to attach to surface. However, the pen-strep treatment can compensate the influence given by PDMS and promote cell spreading (Figure. 2b). Besides influence macrophage sense substrate, it also mediates macrophage-ECM interaction. Type I, IV collagen and laminin are common ECM found in tissue. Pen-strep was found to decrease cell roundness on type I collagen and laminin coated surface (Figure. 2c, f). In contrast, pen-strep increases macrophage roundness on type IV collagen (Figure. 2d). Despite ECM, macrophage sensation to polypeptides which simulate ECM function also tested. Poly amino acid and poly-RGD peptides are widely used in cell biology experiment to help cell attachment [11]. Pen-strep treatment also found to promote cell spread on these substrates coated surface (Figure. 2e, g). These results indicate that pen-strep influence macrophage sense microenvironment factors, including substrate type and ECM coating.

**Figure 2:**
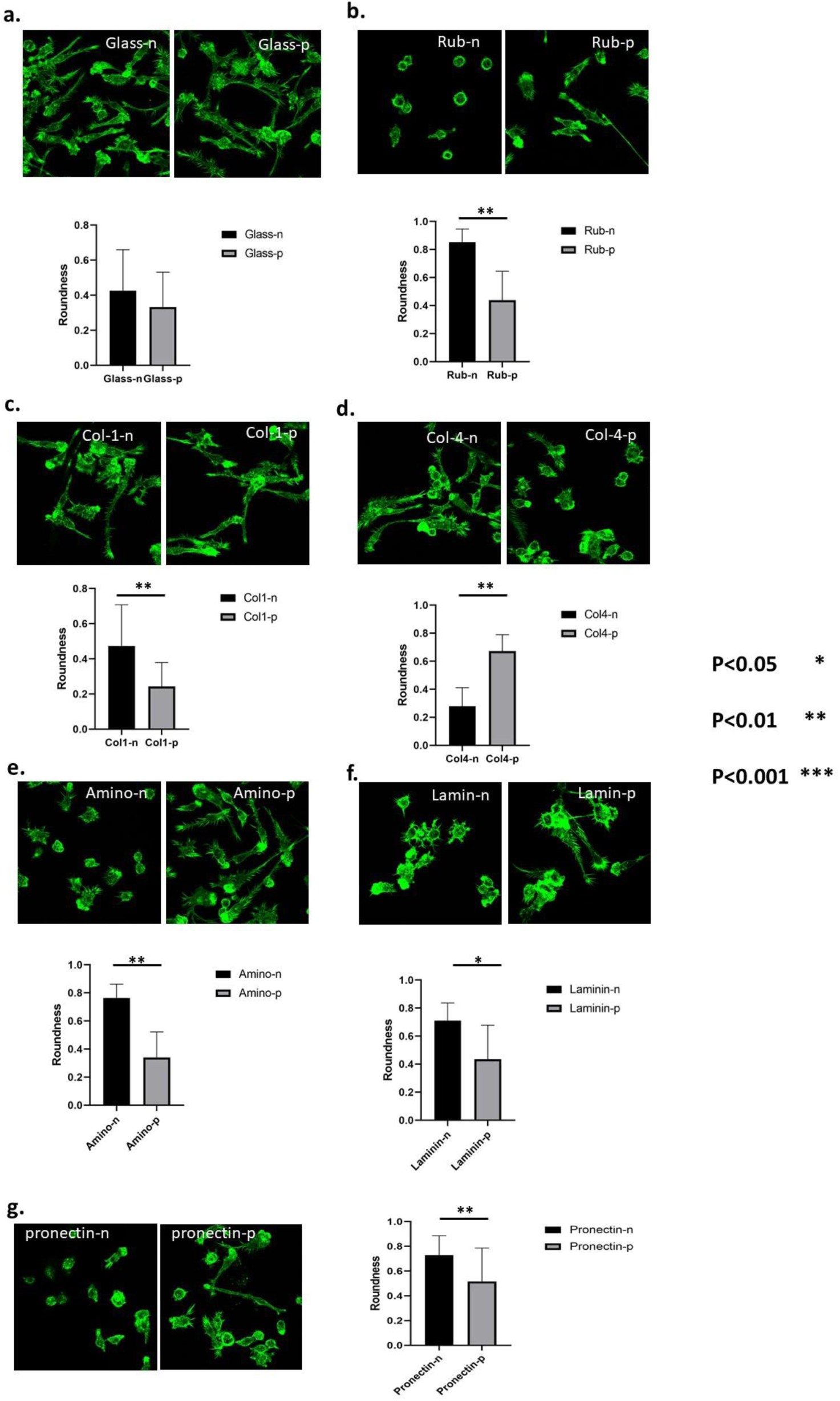
Change of cell morphology and roundness on different substrate and ECM coating: Glass-n: without treatment, Glass-p: Pen-strep treatment a) Glass; b) Rub: PDMS rubber; c) Col-1: Type I collagen; d) Col-4: Type IV collagen; e) Amino: poly amino acid; f) Lamin: Laminin; g) Pronectin: Poly-RGD peptides.

### Pen-strep effect modulus and adhesion gene expression

To investigate the mechanism of these phenomena, the expression transcription factors and surface protein influence cell elasticity and focal adhesion were analyzed. After treating macrophages on glass with pen-strep for 24 hours, gene expression level of TAZ, Egr-1, YAP-1, vinculin, paxillin and β 1 Integrin was evaluated. RT-qPCR results indicate pen-strep treatment upregulated the expression of TAZ and YAP-1 protein (Figure. 3a, c). Furthermore, it also downregulates β 1 integrin which controls cell sensation to the ECM (Figure. 3f). However, pen-strep does not affect paxillin, vinculin and Egr-1 expression.

**Figure 3:**
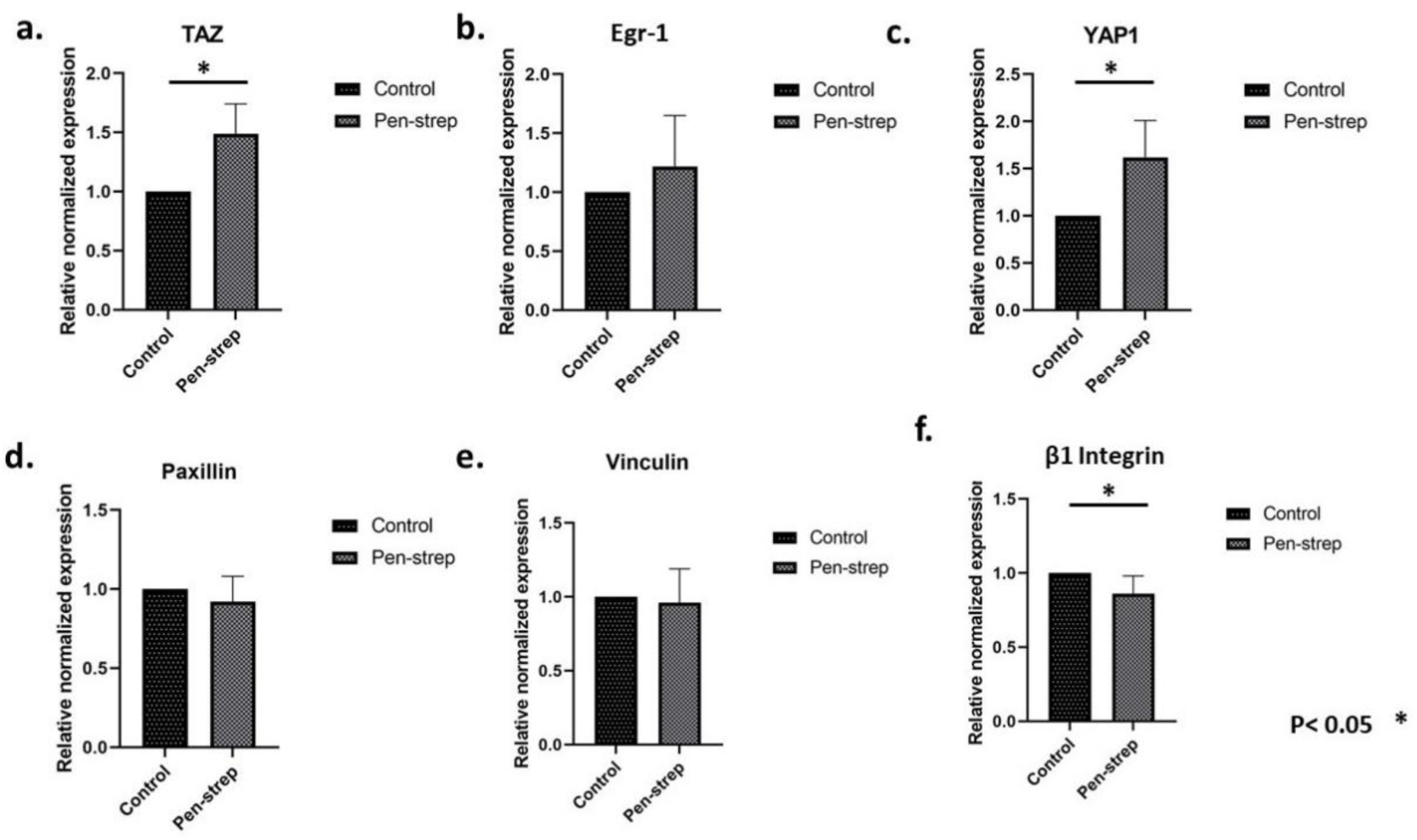
Expression level change of TAZ, Egr-1, YAP1, Paxillin, Vinculin and β1 integrin in macrophage after 1-day treatment. a) TAZ; b) Egr-1; c) YAP-1; d) Paxillin; e) Vinculin; f) β1 integrin.

## Discussion

These data indicate that pen-strep can mediate macrophage modulus, morphology, sensation of ECM substrate and corresponding gene expression. Despite LPS, pen-strep become the other chemicals which can influence macrophage modulus and gene expression pattern. Consequently, the usage of pen-strep in cell culture media for macrophage mechanobiology study become a concern. Also, results imply that pen-strep used in antibiotic treatment may influence macrophage function in inflammation process.

Under pen-strep stimulation, the young modulus of the macrophage on glass increased. Previous research has shown that cell modulus is mediated by cell spread [12]. When macrophage elasticity increase, its roundness also decreased at the same time. This result is consisting with previous research shown that cell with larger spread area has higher elasticity. As cell spread on substrate surface, cell height decreases and cell nuclear elasticity increases. Finally, it increases overall cell modulus [13]. However, roundness of macrophage on glass does not decrease significantly under pen-strep treatment (Figure. 2a). Influence on transcription factor become a possibility to regulate cell plasticity. Further experiments showed that pen-strep treatment also enhances the expression of YAP-1 and TAZ which regulate cell stiffness. YAP-1 and TAZ regulate cell stiffness base on Hippo pathway. Nuclear aggregation of YAP and TAZ can regulates cell modulus [14]. Therefore, the expression and nuclear localization of YAP-1 and TAZ trigger on Hippo pathway and finally mediates cell modulus increase.

In contrast to cell modulus, pen-strep treatment reduced macrophage focal adhesion on day 2 and it increases to normal state after three days of treatment. Usually, cell focal adhesion is regulated by integrin, which cluster can promote phosphorylation of focal adhesion-associated tyrosine kinase [15]. Integrin also can recruit Src kinase and talin to trigger on the corresponding pathway which regulate cell adhesion. In this process, Vinculin control focal adhesion through interacting with talin and actin directly [16]. Similarly, paxillin phosphorylated by Src and interact with actin, further influence cell-ECM attachment [17]. However, after 1-day treatment, both paxillin and vinculin expression did not have significant decrease. Pen-strep only has negative effect on β1 integrin expression, which participated in cell-ECM interaction. Results imply that the reduction of β1 integrin on cell surface may influence focal adhesion on day 2. However, paxillin and vinculin regulated pathway cannot be influenced by pen-strep treatment. Therefore, the influence given by pen-strep is reduced after day 3 treatment. The pathway to give this influence still remain to be investigated.

The cell morphology is influenced by its microenvironment including substrate stiffness and ECM coating on surface [18, 19]. β1 integrin interacts with various ECM and influences cell morphology. Cell spreading was promoted after integrin interact with fibronectin and laminin [15, 20]. However, although β 1 integrin expression level decreased under pen-strep treatment, cell still tends to spread on type I collagen, laminin, poly-RGD peptides and poly amino acid coated surface. The only exception is type IV collagen coated surface, macrophage tends to shrink under pen-strep treatment. It implies that despite integrin, pen-strep trigger on another signal pathway to mediate cell morphology and recognition of ECM. Different combinations of integrin α and β subunits perform different affinity to ECM [21]. Another type of subunits of integrin could be upregulated to compensate the reduce of β1 integrin and increase integrin affinity to ECM. More investigation remains to be done on the mechanism for pen-strep to influence macrophage to interact with ECM.

In summary, we found that pen-strep can increase macrophage modulus and its sensation to microenvironment. Phalloidin staining indicated pen-strep promote cell spread and induce nuclear localization of YAP-1 and TAZ protein. The downregulation of β1 integrin under pen-strep treatment may influence the cell focal adhesion in a short time. However, vinculin and paxillin did not participate in this process. Although β1 integrin expression level decreased, the cell on different ECM coated surface still tends to spread except type IV collagen coated surface. It indicates that pen-strep mediates another pathway to influence cell morphology and compensate the influence given by β1 integrin downregulation. Base on the result of experiments, the application of pen-strep in cell culture media need to be concerned when conducting macrophage research.

## References

1. Mosser, D.M. and J.P. Edwards, Exploring the full spectrum of macrophage activation. Nature Reviews Immunology, 2008. 8(12): p. 958–969.

2. Martinez, F.O., L. Helming, and S. Gordon, Alternative Activation of Macrophages: An Immunologic Functional Perspective. Annual Review of Immunology, 2009. 27(1): p. 451–483.

3. Biswas, S.K. and A. Mantovani, Macrophage plasticity and interaction with lymphocyte subsets: cancer as a paradigm. Nature Immunology, 2010. 11(10): p. 889–896.

4. Mantovani, A., et al., The chemokine system in diverse forms of macrophage activation and polarization. Trends in Immunology, 2004. 25(12): p. 677–686.

5. Patel, N.R., et al., Cell elasticity determines macrophage function. PloS one, 2012. 7(9): p. e41024–e41024.

6. McWhorter, F.Y., C.T. Davis, and W.F. Liu, Physical and mechanical regulation of macrophage phenotype and function. Cellular and molecular life sciences: CMLS, 2015. 72(7): p. 1303–1316.

7. Leporatti, S., et al., Elasticity and adhesion of resting and lipopolysaccharide-stimulated macrophages. 2006. 580(2): p. 450–454.

8. Jain, N., J. Moeller, and V. Vogel, Mechanobiology of Macrophages: How Physical Factors Coregulate Macrophage Plasticity and Phagocytosis. Annual Review of Biomedical Engineering, 2019. 21(1): p. 267–297.

9. Lobanovska, M. and G. Pilla, Penicillin’s Discovery and Antibiotic Resistance: Lessons for the Future? The Yale journal of biology and medicine, 2017. 90(1): p. 135–145.

10. Luzzatto, L., D. Apirion, and D. Schlessinger, Mechanism of action of streptomycin in E. coli: interruption of the ribosome cycle at the initiation of protein synthesis. Proceedings of the National Academy of Sciences of the United States of America, 1968. 60(3): p. 873–880.

11. Studenovská, H., et al., Synthetic poly(amino acid) hydrogels with incorporated celladhesion peptides for tissue engineering. Journal of Tissue Engineering and Regenerative Medicine, 2010. 4(6): p. 454–463.

12. Tee, S.-Y., et al., Cell shape and substrate rigidity both regulate cell stiffness. Biophysical journal, 2011. 100(5): p. L25–L27.

13. Vichare, S., M.M. Inamdar, and S. Sen, Influence of cell spreading and contractility on stiffness measurements using AFM. Soft Matter, 2012. 8(40): p. 10464–10471.

14. Nardone, G., et al., YAP regulates cell mechanics by controlling focal adhesion assembly. Nature Communications, 2017. 8(1): p. 15321.

15. Kornberg, L., et al., Cell adhesion or integrin clustering increases phosphorylation of a focal adhesion-associated tyrosine kinase. 1992. 267(33): p. 23439–42.

16. Humphries, J.D., et al., Vinculin controls focal adhesion formation by direct interactions with talin and actin. The Journal of cell biology, 2007. 179(5): p. 1043–1057.

17. Turner, C.E., Paxillin and focal adhesion signalling. Nature Cell Biology, 2000. 2(12): p. E231–E236.

18. Goodman, S.L., R. Deutzmann, and K. von der Mark, Two distinct cell-binding domains in laminin can independently promote nonneuronal cell adhesion and spreading. The Journal of cell biology, 1987. 105(1): p. 589–598.

19. Yeung, T., et al., Effects of substrate stiffness on cell morphology, cytoskeletal structure, and adhesion. Cell Motility, 2005. 60(1): p. 24–34.

20. Boudreau, N.J. and P.L. Jones, Extracellular matrix and integrin signalling: the shape of things to come. The Biochemical journal, 1999. 339 (Pt 3)(Pt 3): p. 481–488.

21. Anderson, L.R., T.W. Owens, and M.J. Naylor, Structural and mechanical functions of integrins. Biophysical reviews, 2014. 6(2): p. 203–213.

